# Sensitive Transcriptomics and Genotyping reveals function of genetic variants in immunity

**DOI:** 10.1101/2025.03.12.642879

**Authors:** Kai Liu, Anders Rasmussen, Wenkai Han, Qiyu Gong, Seneca Bohley, Dawn Chen, Masahiro Kanai, Regan Maronick, Zhangyuan Yin, Chenlei Hu, Tanvi Jain, Jorge D. Martin-Rufino, Vijay G. Sankaran, Orr Ashenberg, Mark J. Daly, Fei Chen, Daniel B. Graham, Ramnik J. Xavier

## Abstract

Advancements in genomics have revolutionized human genetics by defining the genetic architecture of disease. However, identifying causal variants and their mechanisms of action remains a challenge in translating genetics into therapeutic interventions. Here, we developed Sensitive Transcriptomics And Genotyping by sequencing (STAG-seq), a high-throughput platform designed to define mechanistic genotype-phenotype relationships through simultaneous single-cell measurements of genomic DNA and RNA transcripts. Combined with base-editing, STAG-seq enables functionalization of variants in relevant cellular contexts. We demonstrate the applicability of this approach in several settings. First, we screened genetic perturbations to identify monoallelic and biallelic variant effects in primary human macrophages treated with innate immune stimuli. Next, we phenotyped clinically relevant missense variants associated with immunodeficiency and autoimmunity. Finally, we defined a noncoding variant in a pleiotropic autoimmunity locus that governs TNRC18 expression in primary T cells. STAG-seq thus enables variant phenotyping at scale to advance functional genomics and disease biology.

## Introduction

Understanding the functional impact of DNA variation is a central mission of genetics. While human genetics has rapidly enabled discovery of genetic variation associated with a broad range of diseases, there remains a significant bottleneck in translating these discoveries into identification of causal variants and their mechanisms of action. This is due, in large part, to the need to characterize variant effects within relevant primary cell type contexts, as variant effects are highly context-dependent, influenced by cell type, cell state, and the cellular environment. This complexity often confounds interindividual comparisons in translational studies due to differences in genetic backgrounds and environmental factors^1–3^. To address these challenges, experimental tools that enable the generation of genetic variants within an isogenic background are essential, as they minimize confounding variables and allow for precise comparisons between variants and controls in relevant cellular contexts.

Functional genomics has been revolutionized by high-throughput methods for functional characterization of genetic variants, as well as by genome editing technologies. Deep mutational scanning (DMS) and massively parallel reporter assays (MPRAs) facilitate the simultaneous analysis of a large number of variants, but these approaches are limited in their ability to replicate true allele dosage and endogenous genomic contexts, thereby hindering precise functional characterization of variants^4–7^. CRISPR-based technologies, including base editing and prime editing, enable genetic alterations within the natural genome^8–10^. When integrated with single-cell methods like Perturb-seq, these tools facilitate systematic investigations of variant functions with high-content RNA readouts in pooled screens^11,12^. However, current variant editing tools are often constrained by genomic sequence context, incomplete editing efficiency, and bystander mutations. These limitations obscure the precise determination of edited variants in individual cells, particularly when relying on gRNA proxies and in primary cell contexts where editing efficiencies are low, thereby complicating the accurate interpretation of variant effects.

Overcoming current limitations for variant functionalization will require direct genotyping of variants from the endogenous genome combined with high-sensitivity profiling of the transcriptome within the same single cell. Plate-based methods capable of measuring both DNA and RNA are constrained by low throughput, making them impractical for large-scale studies^13,14^. Microfluidic or split-pool approaches that genotype from open chromatin fragments or transcripts, while scalable, often suffer from high allele dropout rates and may fail to access key coding or noncoding genomic regions, limiting their utility for high-throughput screening^15,16^. Additionally, variants associated with recessive diseases can exert significant phenotypic effects even in a heterozygous state, albeit with subtler signals^17^. Detecting effects of these variants on cellular functions requires methods with allelic sensitivity in genotyping and enhanced power to capture minor transcriptomic changes.

Here, we present Sensitive Transcriptomics and Genotyping by Sequencing (STAG-seq), a droplet-based single-cell method designed to address existing functional genomics challenges. STAG-seq enables the sequencing of hundreds of targeted genomic DNA loci to ascertain genotypes while concurrently profiling the transcriptome with high sensitivity using probe-based detection within the same cell. By integrating STAG-seq with base editing in human primary immune cells, we uncover both mono- and biallelic effects of coding variants implicated in immune-mediated diseases in primary human macrophages. Furthermore, we provide mechanistic insights into a novel noncoding variant within complex autoimmunity locus in primary human T cells, revealing both cis and trans regulatory effects. This approach advances our ability to precisely dissect the functional impact of genetic variants in their native genomic and cellular contexts.

## Results

### STAG-seq enables sensitive profiling of genomic DNA variation and RNA expression in single cells

To effectively capture variants at the genomic DNA level and enhance RNA detection sensitivity, we developed STAG-seq, which integrates a droplet-based single-cell genotyping platform (Tapestri, Mission Bio) with Hybridization of Probes to RNA for sequencing (HyPR-seq)^18^ (Fig. 1A). HyPR-seq offers significant advantages by enabling the design of multiple probes per transcript, thus amplifying the target signal and reducing sequencing costs. Additionally, its DNA probes are compatible with the Tapestri platform’s amplification process.

**Figure 1.**
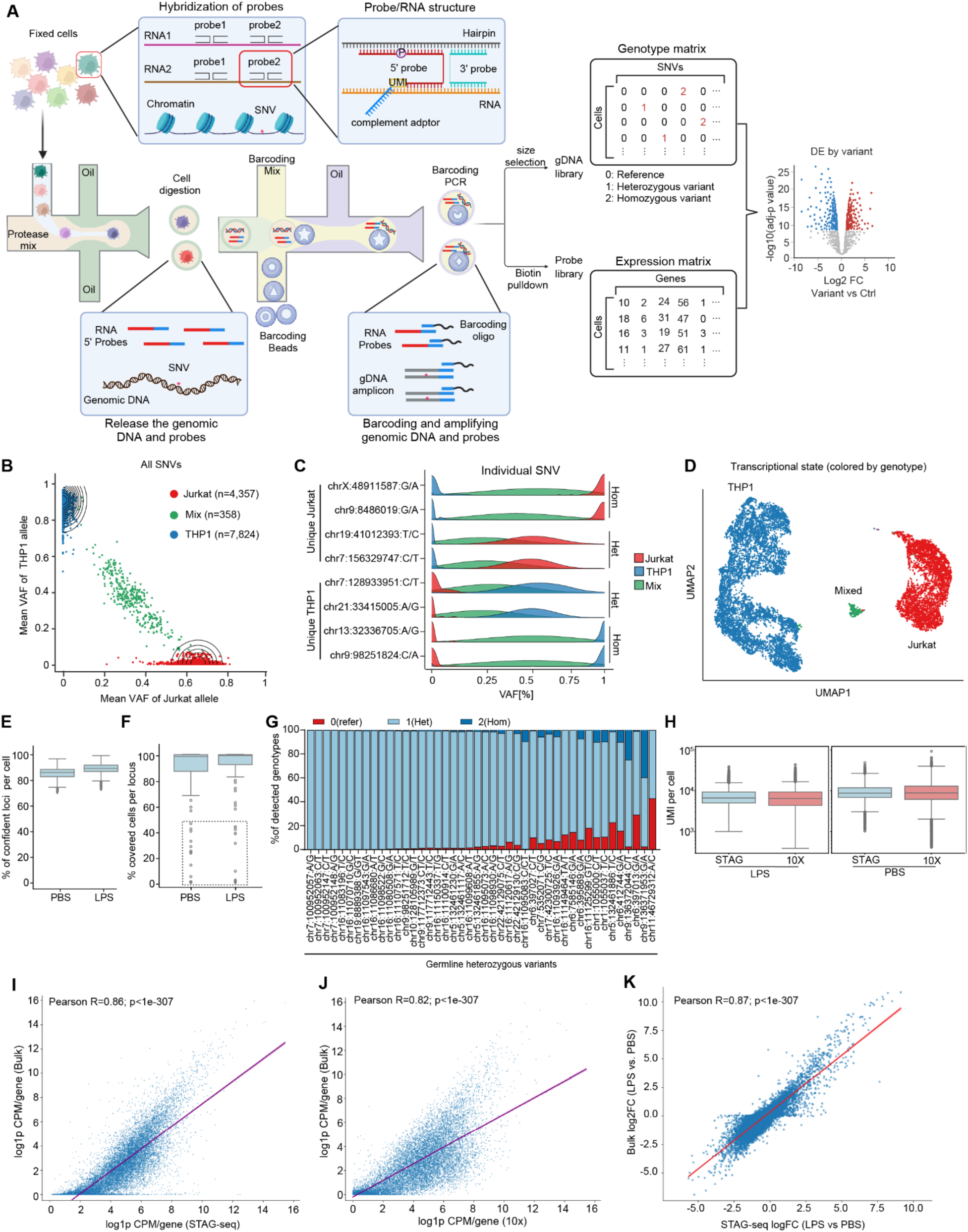
STAG-seq enables single-cell genotyping and parallel transcriptome profiling in high-throughput with sensitivity. (**A**) STAG-seq workflow. (**B**) STAG-seq detection of SNVs from THP1 and Jurkat cells pooled into a heterogeneous population. Called genomic variants were compared with the somatic mutations of each cell type identified by DepMap Omics. Contour lines show cell density across VAF values. (**C**) Distribution of cell populations at selected loci. Cells in (B) projected for each SNV. (**D**) UMAP showing the transcriptional states of pooled THP1 and Jurkat cells from (B), based on the expression of 1,297 genes. Cells are colored by genotype, as in panel (B). (**E**) MDMs were stimulated with LPS or left untreated (PBS) and subjected to STAG-seq with the 50k transcriptome panel and 121-locus gDNA panel. Box plot shows the percentage of high-confidence DNA amplicons in each cell (n= 8,969 and 9,723) for the PBS and LPS samples, respectively. (**F**) Percentage of cells (n= 8,969 and 9,723) that are confidently genotyped for each locus (≥10 reads) for the PBS and LPS samples. The dashed box indicates amplicons that were confidently detected in <50% cells. (**G**) Fractions of genotypes determined by the GATK variant calling for the germline heterozygous SNPs presented in the DNA panel of the PBS sample. Red: GATK called as reference; light blue: GATK called as heterozygous variants; Dark blue: GATK called as homozygous variants. (**H**) Box plots comparing the distribution of total UMIs per cell between STAG-seq and 10x for indicated treatments. (**I**) Correlation of mean CPM/gene between bulk RNA-seq and STAG-seq of 17,885 genes in PBS-treated samples. (**J**) Correlation of mean CPM/gene between 10x and bulk RNA-seq of 17,885 genes in PBS-treated samples. (**K**) Correlation of expression changes of 17,885 genes (LPS vs PBS) between bulk RNA-seq and STAG-seq.

We benchmarked the performance of STAG-seq by evaluating its capability to genotype and phenotype simultaneously in a mixed population of THP1 and Jurkat cells. Accordingly, we designed a custom 2,614-probe panel targeting 1,297 genes and a gDNA amplicon panel detecting 109 distinct chromosomal loci. STAG-seq effectively identified unique single nucleotide variants (SNVs) in both cell lines, separating them by genotype into different populations, and only a small proportion (2.9%) of mixed cell doublets were observed (Fig. 1B). The mean variant allele frequency (VAF) was 0.92 for the homozygous SNVs and 0.53 for the heterozygous SNVs. At each locus, the mixed cells are centered around VAF 0.5 for the reference/homozygous and 0.25 for the reference/heterozygous, indicating equal capture of all alleles (Fig. 1C). To demonstrate the fidelity of concurrent genotyping and phenotyping, we integrated genotype and gene expression profiles within individual THP1 and Jurkat cells. The transcriptional states closely matched genotype identities of the different cell lines, showing a 99.6% overlap (Fig. 1D).

To extend the application of STAG-seq to more complex biological samples, such as tissue analyses, we scaled up by designing an extensive set of 53,540 probe pairs targeting 18,081 genes (referred to as the 50K-set) To assess the genotyping quality and transcriptome fidelity in capturing cell state changes, we applied the 50K-set in conjunction with a DNA panel covering 121 genomic regions to MDMs stimulated with either lipopolysaccharide (LPS) or treated with phosphate buffered saline (PBS) as a control. Concurrently, we conducted bulk DNA sequencing to ascertain the true genotypes, alongside bulk RNA-seq and 10x 3′ scRNA-seq to profile gene expression. With STAG-seq, a median of 86.0% and 89.2% of genomic loci were confidently genotyped per cell in the PBS and LPS samples, respectively (Fig. 1E). While a small fraction of loci (11 out of 121) were genotyped in 5-50% of cells, the remaining amplicons achieved higher detection rates, resulting in a median per-locus detection rate of 99.1% and 98.4% for LPS- and PBS-treated cells, respectively (Fig. 1F). We further evaluated 44 heterozygous germline variants identified by bulk DNA sequencing. Although some loci have a higher allele dropout rate than others, an average of 91.4% genotyping accuracy was achieved, demonstrating the quality of genotyping maintained with scaled transcriptomic measurements (Fig. 1G).

To determine the performance of transcript detection, we subsampled the STAG-seq and 10x sample to have an equal number of total reads (250M) and evaluated the abundance of unique molecular identifiers (UMI) in each cell for the 18,081 genes.

STAG-seq obtained a median of 9,879 and 7,603 UMI per cell for the PBS and LPS sample respectively, showing comparable detection with 10x scRNA-seq (10,435 and 7,255) (Fig. 1H). Gene expression correlation between STAG-seq and bulk RNA-seq was notably strong (Correlation Coefficients = 0.86 (Fig. 1I), comparable to that observed between 10x and bulk RNA-seq (Correlation Coefficients = 0.82 (Fig. 1J). A pseudobulk differential expression analysis further demonstrated that STAG-seq accurately reflected macrophage state changes in response to LPS stimulation with a correlation coefficient of 0.87 (Fig. 1K). Overall, these findings underscore the robustness and sensitivity of STAG-seq in concurrently detecting single-cell DNA and RNA, positioning this technology as a powerful tool for phenotyping genetic variants at scale.

### STAG-seq enables phenotyping variants at scale with genotype by stimulation matrix screens in innate immunity

Innate immune cells, such as macrophages and dendritic cells, express a diverse array of pattern recognition receptors and cytokine receptors, enabling them to respond to specific pathogen threats. Variants associated with innate immunodeficiencies may manifest phenotypes only in particular cell types and under specific biological contexts. We developed STAG-seq with the throughput to support the systematic study of variants in a matrix format wherein multiple variants are individually installed in primary immune cells by base editors and phenotyped across a panel of extrinsic stimuli.

Focussing on a panel of innate immune stimuli, we targeted key genes in pathways downstream of these signals in MDMs and evaluated the ability of STAG-seq to reveal both monoallelic and biallelic variant effects.

First, we optimized base editing in MDMs through electroporation of synthetic sgRNA targeting loci of interest and mRNA encoding various base editors. To expand the range of targetable variants and enhance editing efficiency and precision, we assembled a toolbox to produce base editor mRNAs with combinations of different deaminase variants (evoAPOBEC1, BE4Max, ABE8e and TadCBE) with PAM-flexible SpCas9 variants (NG and SpRY), as well as introducing the narrow-window mutations into the deaminases^19,20^. By targeting sites with NGG or NRY PAMs, we found that evoAPOBEC1(YE1)-SpRY, TadCBEd(V106W)-SpRY and ABE8e(F148A)-SpRY achieved high efficiencies (Fig. 2B).

**Figure 2.**
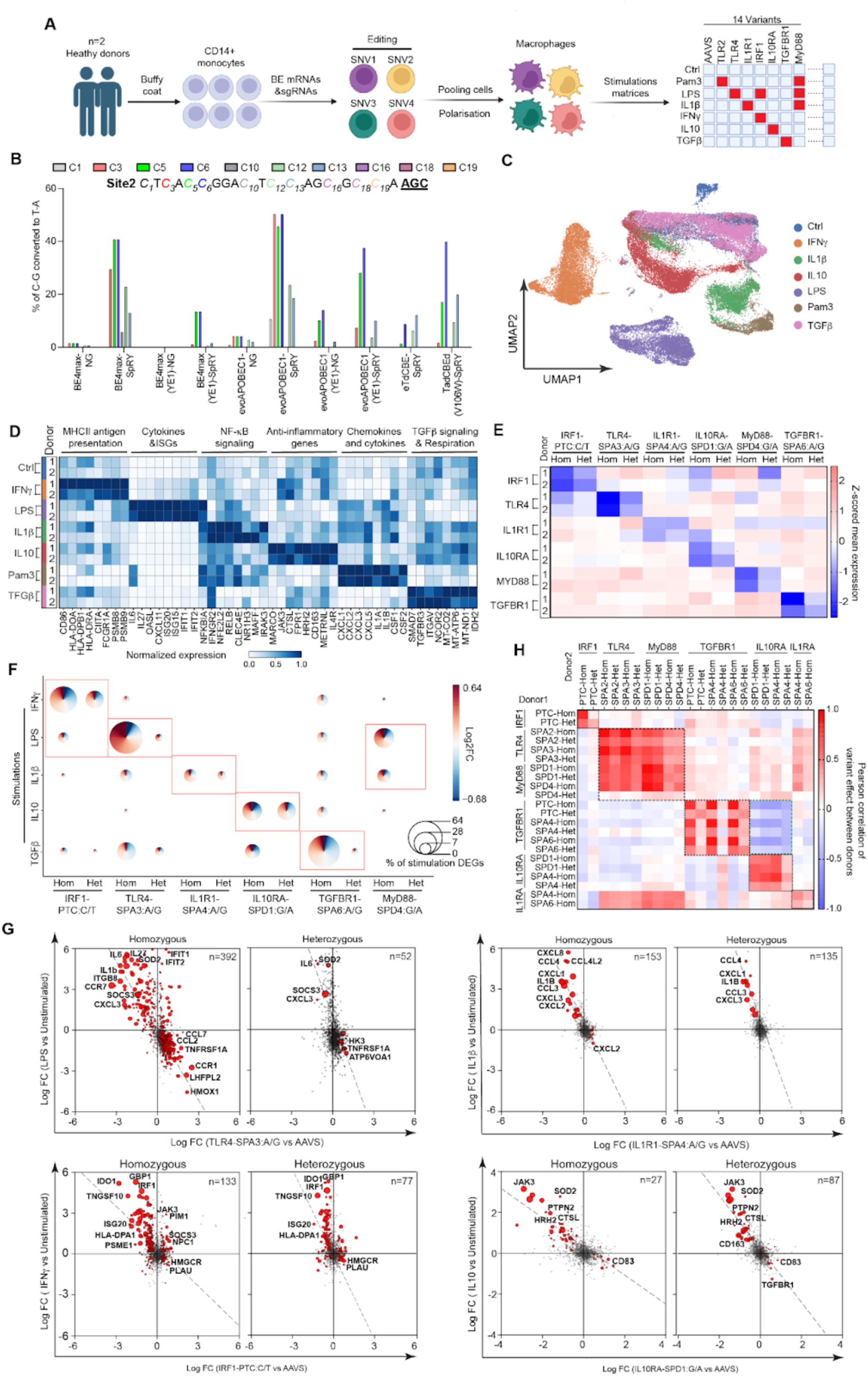
STAG-seq deciphers the function of single base changes in a matrixed base editor screen. (**A**) Schematic workflow of the matrix screen with 14 splice-site or nonsense mutations of 8 genes under 7 stimulation conditions. The red box in the matrix signifies expected variant responses with the given stimuli. (**B**) Base editor precision and efficiency using multiple combinations of CAS9-cytosine deaminase enzymes with a control guide RNA targeting a locus with NRY PAMs.. (**C**) UMAP plot showing STAG-seq transcriptome with 1,275 genes in 56,783 cells. Edited MDMs were stimulated with the indicated TLR ligands or cytokines. (**D**) Heatmaps showing the standardized mean expression of marker genes (X-axis) in the unedited cells of two donors after indicated stimulation (Y-axis). Columns represent individual genes with mean expression. (**E**) Heat maps showing the effect of variant alleles (columns) on gene expression (rows) in cis at steady state (rows are Z-scored). Base changes are indicated after the colon. (**F**) Bubble pie plot showing the specificity of variant allele effects after stimulation. Bubble size indicates percentage of stimulation-induced differentially expressed genes (DEGs) in unedited cells observed as DEGs in variant vs AAVS under the same stimulus. For stimulation DEGs, adjusted P<0.0001, logFC >0.5 or <-0.5. For variant DEGs, adjusted P<0.05. Het, heterozygous; Hom, homozygous. Base changes indicated after the colon. Red boxes indicate expected responses. (**G**) Quadrant plots showing variant allele effects on expression of all measured genes under indicated stimulations. X-axis: Log FC of gene expression between variant cells and AAVS-edited cells under indicated stimulations in Y-axis. Red dots indicate genes with significant changes in variant cells. Dot size, -Log10 of the adjusted P value. (**H**) Heatmap showing the Pearson correlation coefficient of variant allele effects on the relevant stimulation responsive genes (LogFC>0.5 or -0.5<, adjustP<1-e30) between two donors. *IRF1* variants under IFNγ stimulation; *TLR4* and *Myd88* variants under LPS stimulation; *TGFBR1* variants under TGFβ stimulation; *IL10RA* variants under IL-10 stimulation; *IL1RA* variants under IL-1β stimulation. Dashed boxes indicate groups of variants expected to affect the same programs.

We next conducted a “matrix screen” in MDMs isolated from two donors, treating the cells with various stimuli (PBS, LPS, IFN-γ, Pam3CSK4, IL1β, IL10 and TGFβ) and applied 2,740 probes targeting 1,275 genes involved in key immune pathways as transcriptomic readouts. Using base editors, we introduced 14 nonsense or splice site mutations in receptors and key signaling proteins downstream of these stimuli (TLR2, TLR4, MyD88, IRF1, IL1R1, IL10RA, and TGFBR1), along with on-target control edits in the safe harbor AAVS1 locus. Pooling all variant cells for each stimulation allowed us to examine the specific responses of variant-stimulation pairs (Fig. 2A). The stimulations induced distinct gene expression programs that were shared by both donors (Fig. 2C). Furthermore, STAG-seq discerned the unique response signatures for each stimulus: IFN-γ upregulated antigen presentation-related genes; LPS and Pam3CSK4 predominantly induced proinflammatory cytokine production; IL1β triggered NF-kB signaling; IL-10 specifically upregulated anti-inflammatory genes; and TGFβ primarily impacted TGFβ signaling and cell metabolism characteristic of alternative macrophage activation (Fig. 2D).

To assess the functional impact of edited variants, we first examined effects on expression of their respective target genes. Homozygous mutations generally reduced target gene expression, while heterozygous mutations showed subtler but significant decreases, demonstrating the ability of STAG-seq to detect and distinguish *cis* effects of homozygous and heterozygous variants (Fig. 2E). This is also consistent with the fact that nonsense or splice-site mutations usually trigger the nonsense-mediated mRNA decay (NMD) pathway to destabilize the mutated mRNA. We then investigated *trans*-regulatory effects by measuring the proportion of stimulation-responsive genes affected by each variant. Homozygous receptor variants (TLR4, IL1R1, IL10RA, and TGFBR1) dramatically altered responses to their respective ligands (LPS, IL-1β, IL-10, and TGFβ) with minimal effects from other stimuli, while heterozygous variants exhibited similar patterns but with reduced magnitude (Fig. 2F). We also identified variants with pleiotropic effects. For example, the IRF1 homozygous variant strongly impacted the IFN-γ signaling response and weakly but robustly impacted the LPS response (40.6% of IFN-γ signaling, 5.6% of LPS). Similarly, Myd88 variants altered both LPS and IL-1β responses (22.8% of LPS, 9.7% of IL1β) (Fig. 2F). These results align with previous reports that IRF1 acts as a pioneer transcription factor mediating signaling from LPS and IFN-γ, while MyD88 serves as an essential adapter for both TLRs and IL-1β receptors^21–24^.

To further characterize how edited variants influence stimulation-induced gene expression, we analyzed specific gene sets. Homozygous *TLR4* variants corresponding to nonsense mutations or splice site disruption dramatically reduced LPS-induced cytokines and chemokines (*IL6*, *IL27*, *IL1β*, and *CXCL3*) while increasing LPS-suppressed genes (*CCR1*, *HMOX1*, *LHFPL2*, *TNFRSF1*, and *CLEC7A*), indicating a strong suppression of LPS responses in variant cells (Fig. 2G). The corresponding heterozygous variants showed partial blockade with weaker reduction of *IL6* and *CXCL3* expression (Fig. 2G). Similarly, homozygous variants in *IRF1, IL1R1,* and *IL10RA* strongly inhibited gene expression responses to IFNγ, IL-1β, IL-10, and TGFβ treatment, respectively, with heterozygous variants exhibiting the same trends but with smaller magnitude changes and fewer differentially expressed genes (DEGs) (Fig. 2G). Importantly, the effects of homozygous and heterozygous variants were consistent across different variants of the same gene, different genes within the same pathway (e.g., TLR4 and MyD88), and the same gene in two independent donors (Fig. 2H). Taken together, STAG-seq enabled the systematic identification of mono- and biallelic variant functions in primary human immune cells under diverse biological stimuli, providing a powerful platform for high-throughput matrix screens in functional genomics.

### STAG-seq discerns function for clinically relevant coding variants impacting core immune pathways

We next sought to deploy STAG-seq to reveal the functions of clinically relevant coding variants, aiming to accelerate mechanistic understanding of genetic variants of uncertain significance (VUS). Hundreds of variants have been identified to date that impact different components of the interferon gamma (IFN-γ) signaling pathway through distinct mechanisms of action. Given its central role in autoimmunity, infectious disease, and cancer, we selected variants associated with a variety of disease phenotypes that affect key IFN-γ pathway components—including IFNGR1, IFNGR2, JAK1, JAK2, STAT1, and IRF1—from OMIM and ClinVar (Fig. 3A). We further prioritized 24 Clinvar and 6 control variants across the 6 genes, achieving ≥10% editing efficiency in MDMs isolated from 2 donors for profiling with STAG-seq. Overall, editing efficiencies varied by site, as did bystander editing rates, which ranged from 12% to 88% at different loci, supporting the necessity of precise genotyping to ascertain true variant function (Fig. 3B). After editing, we then pooled all the cells together and treated them with PBS or IFN-γ. With unsupervised clustering by the expression of 1,275 genes (2,740 probes), IFN-γ stimulated cells largely separated from PBS treated cells (Fig. 3C). Notably, two positive control variants at the splice sites of *IFNGR1* and *IFNGR2* nearly abolished cellular response to IFN-γ (Fig. 3D). Even with only five homozygous *IFNGR1* control variant (c.88-2A>G) cells, STAG-seq detected statistically significant inhibition of IFN-γ pathway genes (Fig. 3D).

**Figure 3.**
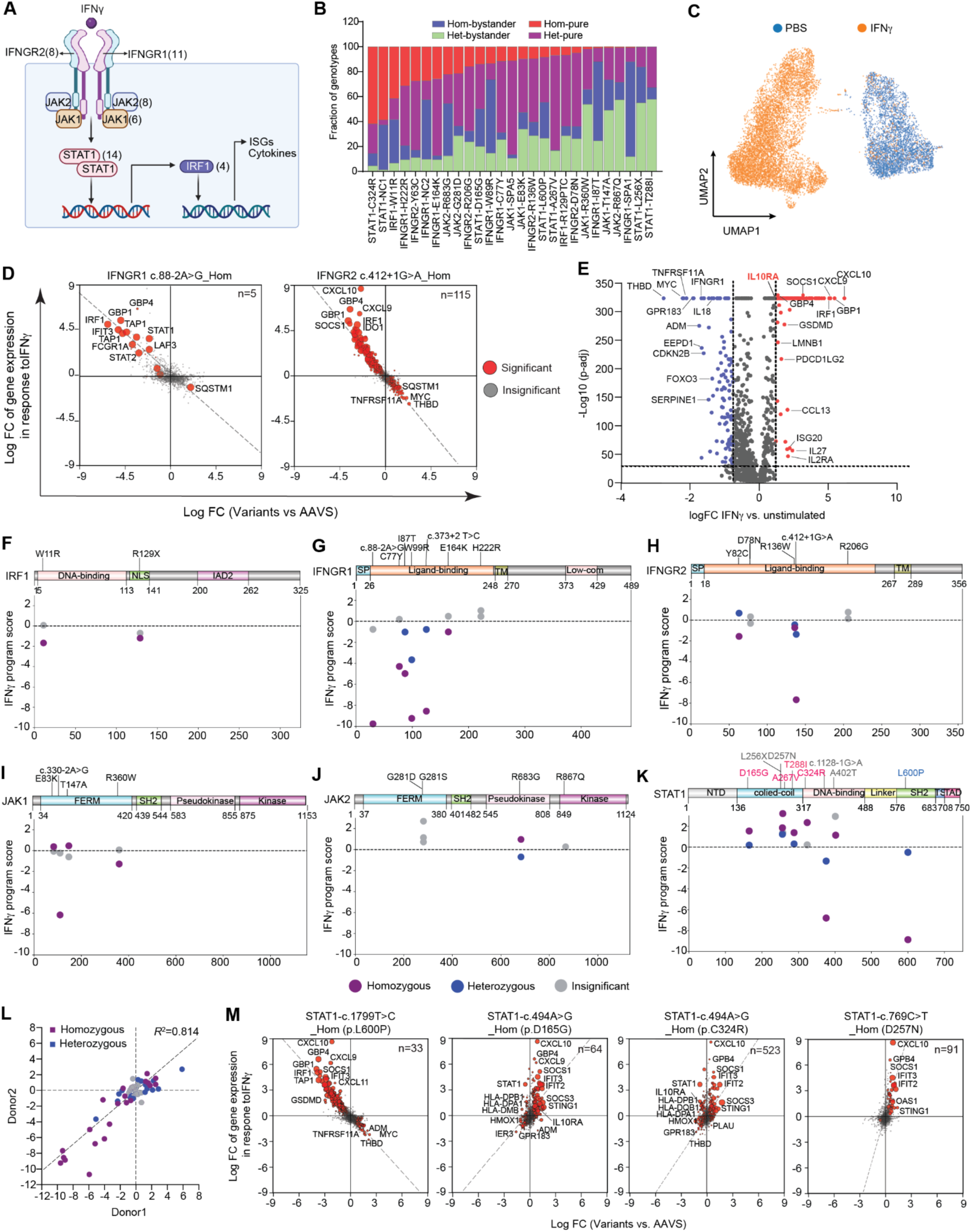
STAG-seq reveals the function of clinically-relevant coding variants in the IFN-γ pathway. (**A**) Schematic of IFN-γ pathway. The number of variants investigated in MDMs for each gene is listed in parentheses. Variants were selected from OMIM and ClinVar based on associations with immune-mediated diseases or cancer. (**B**) Precision and efficiency of editing after variant calling from STAG-seq. Red and purple shows cells containing variants without any other bystander mutations within the 20bp window encompassing the targeted variant. Blue and green shows cells containing variants along with any other mutations (homozygous or heterozygous) within the 20bp target window. (**C**) UMAP plot showing STAG-seq transcriptome of 13,749 unedited cells with detection of 1,275 genes under IFN-γ stimulation. MDMs were stimulated with PBS or IFN-γ. Cells were fixed and hybridized with 2,740 probes for 1,275 genes and then processed into the STAG-seq pipeline with detection of 85 genomic regions. (**D**) Quadrant plots showing the effect of two homozygous IFN-γ receptor variants on all measured genes in response to IFN-γ. Red dots indicate genes with significant changes in variant cells. Dot size, log100 of adjusted P value. (**E**) Volcano plot of IFN-γ response genes in unedited MDMs. Dashed lines indicate the threshold used to select genes (top 100 upregulated and downregulated genes with adjusted P<e30) for calculating gene program scores. (**F-K**) The effect of clinical variants on the IFN-γ program. Positive scores represent an increased response to IFN-γ in edited cells (gain-of-function) while negative scores represent a decreased response to IFN-γ in edited cells (loss-of-function). Purple and blue dots represent pure homozygous or heterozygous, respectively. (**L**) Pearson correlation of the variant allele effects on the IFN-γ program scores in two donors. Each dot represents a variant. P <0.0001, two-tailed t test. (**M**) Quadrant plots showing the effect of *STAT1* variants on all measured genes in response to IFN-γ. “n” indicates the cell number of each variant genotype.

To directly compare the phenotypes and effect sizes of genetic variants, we derived IFN-γ response scores that summarize the transcriptional responses to IFN-γ for each “pure” homozygous and heterozygous variant lacking bystander mutations. Accordingly, negative scores indicate impaired IFN-γ response, whereas positive scores reflect enhanced signaling (Fig. 3, E-K). With this approach, individual variant phenotypes were remarkably consistent across donors, underscoring the robustness of STAG-seq for functional profiling (Fig. 3L). Importantly, the variant effects aligned with the clinical reports. For instance, the immunodeficiency-associated STAT1 variant rs137852678 (L600P) exhibited complete inhibition of IFN-γ signaling in homozygous cells, aligned with its clinical classification as autosomal recessive complete deficiency (Fig. 3K, 3M)^25^.

Having established a framework for accurately genotyping edited variants, we integrated molecular and clinical phenotypes. Variants of *IRF1*, *IFNGR1*, *IFNGR2*, *JAK1*, and *STAT1* that are associated with immunodeficiencies and susceptibility to viral and mycobacterial infections exhibited loss-of-function (LOF) phenotypes and reduced IFN-γ response scores (Fig. 3F-K). In addition to variants linked to infectious disease, we also characterized cancer-associated variants in the IFN pathway, including *JAK2* variant rs1057519721 (pR683G), which has been implicated in pediatric acute lymphoblastic leukemia (ALL)^26–29^. STAG-seq demonstrated an allele-dose effect for this variant, where homozygous cells exhibited a gain-of-function (GOF) phenotype, whereas heterozygous cells displayed a LOF phenotype, (Fig. 3J). *STAT1* variants exhibited the most diverse range of disease associations and functional effects based on IFN-γ responses (Fig. 3K). As a well-known nonredundant signal transducer and transcription activator of types I, II, and III IFN as well as IL-27^30^ pathways, STAT1 mutations underlie diverse clinical phenotypes. The variants classified as complete LOF by STAG-seq are associated with severe immunodeficiencies and susceptibility to viral and mycobacterial infections^31,32^. In striking contrast, variants classified as GOF based on IFN-γ response scores are associated with autoimmunity, and paradoxically, susceptibility to fungal infection^31,33^. These findings are consistent with recent work showing increased Th1 responses underlying impaired antifungal immunity^34^. STAG-seq identified and corroborated additional GOF *STAT1* variants associated with autoimmunity and fungal infection, including rs1574653439 (pC324R), rs1693751220 (pT288I), rs387906759 (pA267V), rs387906764 (pD165G), and also one variant (pD257N) that was identified in a CRISPR screen (Fig. 3, K and M)^31,35^. Notably, although these GOF variants enhanced the expression of most IFN-γ-induced genes (*CXCL10*, *GBP4*, *SOCS1*, *IFIT2*, *SOCS3* and *STING1*), some of them inhibited STAT1 and MHC-II expression after IFN-γ stimulation, indicating a more complex phenotype (Fig. 3M). Additionally, different GOF variants exhibited distinct transcriptional profiles at steady state and after stimulation. While IFN-γ stimulation potentiated the expression of *IL10RA* in control (AAVS1 edited) cells, this response was attenuated by rs1574653439 (pC324R) and enhanced by rs387906764 (pD165G) (Fig. 3E), suggesting distinct mechanisms of action between GOF variants. Together, these findings demonstrate STAG-seq’s ability to systematically resolve both LOF and GOF phenotypes across a spectrum of effect sizes, linking genetic variation to immune dysfunction and disease risk. Finally, to further validate our findings, we compared STAG-seq-discovered phenotypes with ClinVar classifications. All 14 ClinVar-classified pathogenic or likely pathogenic variants exhibited significant IFN-γ response scores, consistent with their known disease associations. Notably, 5 out of 10 variants of uncertain significance (VUS) also displayed significant IFN-γ response scores, indicating that these variants may indeed affect gene function. These results highlight STAG-seq’s potential for reclassifying VUS, offering a powerful framework to bridge genetic variation with functional consequences at single-cell resolution.

### STAG-seq enables functional dissection of noncoding variants within a complex locus linked to autoimmune disease

Population genetics and GWAS predominantly identify noncoding variants with moderate effect sizes on gene expression, and mapping noncoding variants to functional effects on neighboring genes is complicated by cell type specificity and availability of samples with desired variants. This is the case for rs748670681 (chr7:5397122 C/T), a Finnish-enriched noncoding variant on chromosome 7 that exhibits a unique profile of disease associations (MAF = 3.6% in FinnGen; 114-fold Finnish enrichment compared to non-Finnish Europeans). The minor allele imparts risk for IBD, ankylosing spondylitis, and psoriasis (OR = 2.2, 2.3, and 1.5, respectively); but is protective for T1D, multiple sclerosis, Grave’s disease, and autoimmune hypothyroidism^36^ (OR = 0.6, 0.6, 0.5, and 0.9, respectively; Fig. 4A). The locus (800 kb) harboring rs748670681 contains several protein-coding genes (Fig. 4B). Although the variant resides within an intron of *TNRC18*, any gene in the locus could be a cis eQTL target, necessitating empirical evaluation using STAG-seq.

**Figure 4.**
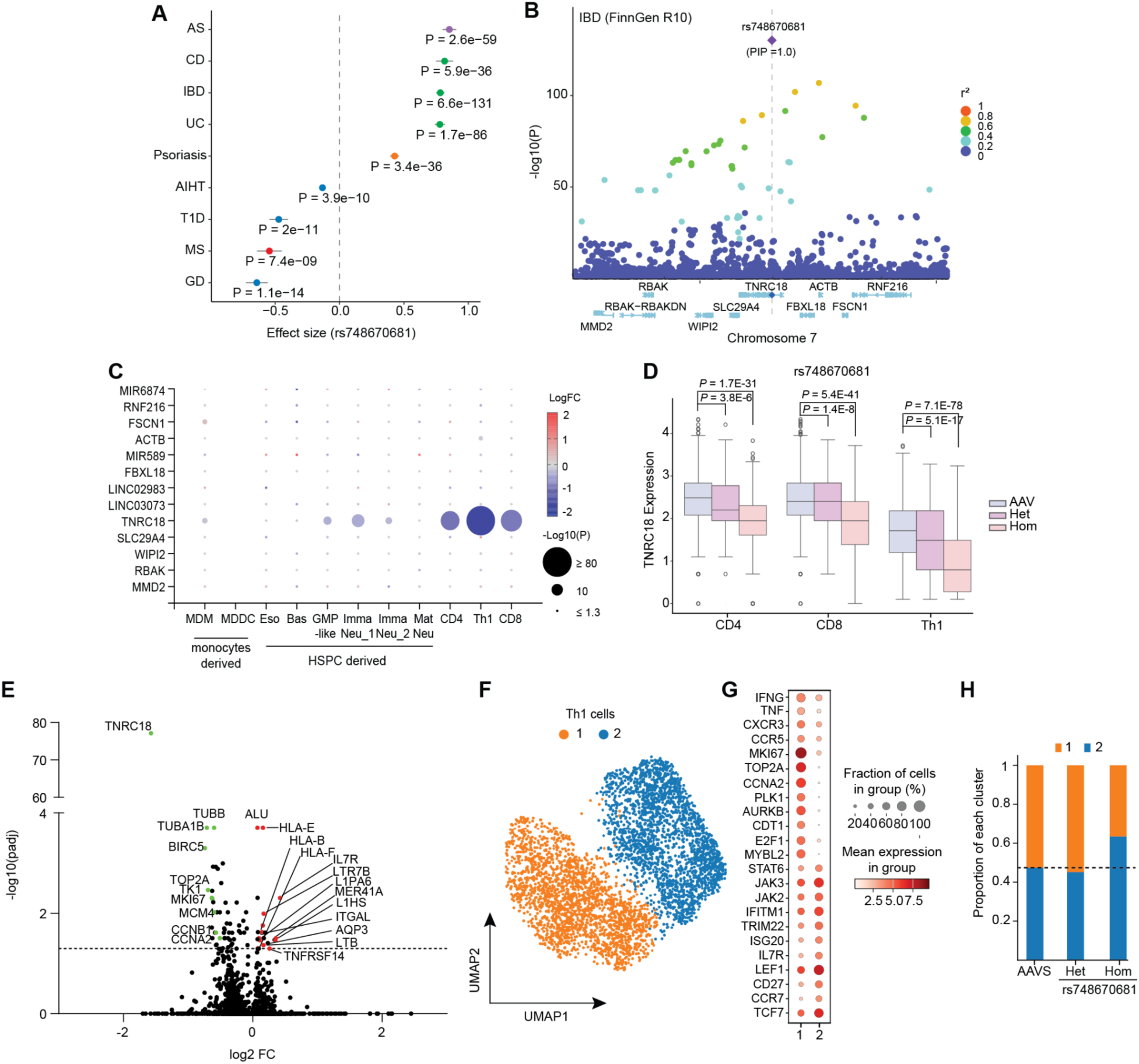
STAG-seq identifies the cis- and trans-effects of a pleiotropic variant linked to autoimmunity. (**A**) Forest plot showing associations of rs748670681 with different autoimmune diseases (Y-axis) in FinnGen R10. P-values were computed using REGENIE^61^. Error bar represents standard error of effect size *β* (log odds ratio). AS, ankylosing spondylitis; CD, Crohn’s disease; IBD, inflammatory bowel disease; UC, ulcerative colitis; AIHT, autoimmune hypothyroidism; T1D, type I diabetes; MS, multiple sclerosis; GD, Grave’s disease. (**B**) Manhattan plot of IBD GWAS data. Fine-mapping identified rs748670681 as an IBD risk factor (posterior inclusion probability [PIP] = 1.0). Color corresponds to a *r*^2^ value to the lead variant rs748670681. (**C**) Mapping cell type specific gene regulation by rs748670681. Dot plot shows the expression changes of genes within the locus in different cell types with the rs748670681 homozygous variant relative to AAVS-edited cells. (**D**) Box plot depicting the cis-regulatory effect of rs748670681 on *TNRC18* expression in CD4, CD8, and Th1 cells. (**E**) Volcano plot illustrating the trans-regulatory impact of rs748670681 homozygous variant determined by STAG-seq. Red dots: highlighted upregulated genes; green dots: highlighted down regulated genes. (**F**) UMAP plot showing 4,354 Th1 cells segregated into two distinct clusters with STAG-seq detection of 2,315 genes. (**G**) Dot plot displaying differentially expressed genes across the clusters identified in (F). (**H**) Cluster composition plot comparing three indicated genotypes. The dash line indicates the proportion of cell clusters in AAVS control.

Given its strong autoimmune association, we introduced the variant into human immune cells of both myeloid and lymphoid lineages. These include monocyte derived macrophages or dendritic cells (MDDCs), granulocytes differentiated from human hematopoietic stem and progenitor cells (HSPCs), activated CD4 and CD8 T cells, as well as ex vivo differentiated Th1 cells Comparing the variant cells to the AAVS1 edited cells, we first examined cis effects of rs748670681 on all genes within the locus and across different immune cell types. The homozygous variant significantly downregulated *TNRC18* expression in specific cell types without impacting other genes (Fig. 4C). A mild cis eQTL effect was observed in granulocyte-macrophage progenitor (GMP)-like cells and immature neutrophils, but not in other myeloid cell types (Fig. 4C). Notably, the cis-eQTL effect was most pronounced in T cells, particularly in Th1 cells, where homozygous variant cells exhibited stronger downregulation of TNRC18 than heterozygous cells, confirming both monoallelic and biallelic cis-eQTL effects (Fig. 4D).

TNRC18 has recently been identified as an epigenetic reader of H3K9me3, enforcing silencing of endogenous retrotransposons in multiple cancer cell lines, although its functions in primary immune cells are still unclear^37^. To examine potential trans effects of rs748670681, we designed 1,234 probes targeting 224 retrotransposons at family or subfamily levels and 4,605 probes targeting 2,021 coding genes. Among all tested cell types, rs748670681 showed the strongest cis effect on *TNRC18* expression in Th1 cells, with broad reactivation of ERVs, human-specific L1HS (LINE-1 Class) and the Alu element (SINE class) (Fig. 4E). The variant cells also upregulated expression of IL7R, MHCI molecules and TNF superfamily members (*TNFRSF14*, *LTB*), while downregulating cell cycle genes, suggesting a shift from proliferation to a survival/memory-like cell state (Fig. 4E)^38–42^.

To further evaluate the impact of rs748670681 on cell state, we investigated the heterogeneity of Th1 cells, which segregated into two major clusters (Fig. 4F). Cluster 1 cells expressed Th1 cytokines and chemokines (*IFNG*, *TNF*, *CXCR3*) and show a clear feature of cell cycle progression and mitosis, suggesting they are activated and proliferating cells (Fig. 4G). Cluster 2 cells expressed higher levels of stemness/memory genes (*IL7R*, *TCF7*, *LEF1*, *CD27*) and tissue homing markers (*CCR7 CXCR6*), suggesting that they are memory-like cells that transiently enter into quiescent state for long-term survival (Fig. 4G). Those cells also exhibit induction of ISGs (Fig. 4G). Given the fact that Th1 cells are a major source of IFN-γ and respond to it through positive feedback, cluster 2 cells might be primed for inflammatory responses. This is also consistent with a previous report that TEs contribute to the expression of ISGs by facilitating the binding of IFN-γ induced transcription factors^43^. Compared to the AAVS edited control, rs748670681 homozygous variant leads to a significantly increased proportion of memory-like cells (Cluster 2, χ² = 44.14, p < 0.0001), indicating a variant-driven shift toward long-live memory states, which may contribute to tissue inflammation (Fig. 4H). Together, our findings revealed a cell type specific cis- and trans-effect of the autoimmunity associated variant rs748670681, and also provide mechanistic insights into its potential role in autoimmune disease. Thus, by integrating genome editing, single-cell genotyping and transcriptomic profiling across diverse immune cell states, STAG-seq uncovered a functional mechanism for a rare noncoding variant in primary immune cells, providing a powerful framework for dissecting the molecular basis of autoimmune risk variants.

## Discussion

Understanding how genetic variation - germline or somatic - contributes to disease remains a central challenge in human genetics. While genome-wide association studies (GWAS) and clinical sequencing have identified thousands of disease-associated variants, the vast majority remain functionally uncharacterized, particularly those with subtle regulatory effects or that function in a context-dependent manner. Furthermore, the heterozygous nature of most variants in the human genome underscores the critical need to resolve allele-specific contributions to disease phenotypes. To address those challenges, we developed STAG-seq, a droplet-based method for single-cell gDNA amplicon sequencing combined with targeted transcriptomics. STAG-seq enables parallel genotyping of hundreds of loci and profiling of thousands to tens of thousands of genes across 10⁴–10⁵ cells in a single experiment. This approach offers three key advancements: 1) high-sensitivity genotyping with minimal allelic dropout (≤2.5%), enabling precise detection of both homozygous and heterozygous variants at scale; 2) enhanced RNA detection with better quantitative accuracy than standard 3’ transcriptome scRNA-seq, empowering functional studies of variants with subtle effect sizes; 3) flexible targeting of genomic regions and RNA transcripts, allowing researchers to focus on specific biological questions while reducing sequencing costs. Additionally, our in-house protocol for synthesizing customized probe pools simplifies adoption of this technology.

The sensitivity of STAG-seq arises from its direct, accurate gDNA sequencing with exceptionally low allelic dropout rates, enabling experiments that are infeasible with existing methods. By comparison, Perturb-seq approaches rely on detection of sgRNA in edited cells based on the prior that cells expressing these guides will have a predictable editing outcome^11,12^. However, editing efficiencies vary by locus^44^, and bystander editing is common. Proxy-based genotyping approaches utilize a transcribable decoy genomic target sequence to detect edits and infer editing at the endogenous genomic locus^45–47^. This genotype inference is an improvement, but fails to directly reveal variant zygosity in editing endogenous loci and stochastic bystander editing. Alternatively, direct genotyping from the transcriptome or chromatin fragments enables detection of potential edits, but is limited by cell dropouts, access to the region of interest, and gene expression levels^15,16,48^. Paired genome and transcriptome sequencing approaches represent an advancement towards improved genotyping and coupling to single cell gene expression profiles^13,49–54^, although these are largely constrained to the throughput of arrayed plate-based formats. In contrast, STAG-seq offers orders of magnitude higher throughput with added sensitivity for paired genotyping and gene expression profiling, which could further be used to validate and improve AI models for genomics^55–57^.

By integrating base editing with single-cell multi-omics, STAG-seq addresses a significant obstacle in translating discoveries from human genetics into mechanistic insights of disease. STAG-seq’s ability to accurately genotype installed edits while sensitively capturing transcriptomic changes in the same single cell unlocks systematic exploration of genotype-phenotype relationships across diverse disease contexts.

## Materials and Methods

### Primary human cells

Primary human CD34+ hematopoietic stem and progenitor cells (HSPCs) from mobilized peripheral blood of healthy donors were purchased from the Fred Hutchinson Cancer Research Center. Monocytes and T cells were isolated from buffy coats obtained from de-identified donors obtained from Research Blood Components (Watertown, MA).

### Human primary monocyte preparations

Monocytes were isolated from buffy coats using human Monocyte Enrichment Cocktail (StemCell Technologies, 15068) with Sepmate tubes (StemCell Technologies, 85460) and ACK lysis buffer (ThermoFisher, A1049201) following the manufacturer’s protocols. Monocytes were either edited directly by nucleofection or aliquoted and frozen at -80°C in FBS with 10% DMSO for later use.

### Human primary T cell preparations

PBMCs were first isolated by density gradient centrifugation with Ficoll-Paque™ (Cytiva, 17144003) and ACK lysis buffer (ThermoFisher, A1049201) following the manufacturer’s protocols. Then CD4 or CD8 positive T cells were isolated from PBMCs using EasySep™ Human CD4+ or CD8+ T Cell Isolation Kit (StemCell Technologies, 15068) following the manufacturer’s protocols. T cells were either activated directly or aliquoted and frozen at -80°C in FBS with 10% DMSO for later use.

### Cell cultures and differentiation

#### Primary Monocyte-derived macrophage cultures

Monocyte derived macrophages (MDMs) were cultured at a density of 5×10^5^/ml in macrophage medium (DMEM high glucose supplemented with 10% FBS [Sigma, F2442], 1% GlutaMAX [Life Technologies, 35050079], 1% penicillin/streptomycin [Life Technologies, 10378016], and M-CSF 100 ng/ml [Peprotech, 315-02]). Medium was changed every 2 days by replacing half of the volume with fresh medium.

After 7 days of differentiation, MDMs were collected by TrypLE (ThermoFisher,12604039) and replated with fresh medium. After 24 hours of recovery, the cells were stimulated with TLR ligands or cytokines as follows: Pam3CSK4 (100 ng/ml, Invivogen, tlrl-pms) for 4 hours; LPS (10ng/ml, Invivogen, tlrl-pb5Ips) for 4 hours; IFN-γ (10 ng/ml, Peprotech, 300-02) for 6 hours, TGF-β1 (100 ng/ml, Peprotech, 100-21) for 6 hours; IL-1β (100 ng/ml, Peprotech, 200-01B) for 6 hours; IL10 (100 ng/ml, Peprotech, 200-10) for 6 hours. The cells were harvested with TrypLE for STAG-seq.

### HSPC myeloid differentiation

Primary human CD34+ hematopoietic stem and progenitor cells (HSPCs) from mobilized peripheral blood of healthy donors were purchased from the Fred Hutchinson Cancer Research Center. The cells were seeded in 6-well plates at a density of 1×10^5^/ml in StemSpanTM SFEM II medium (stemCell Technologies, 02690) with 1% L-glutamine (Thermo Fisher Scientific, 25-030-081) and 1% penicillin/streptomycin (Life Technologies, 10378016) and also cytokine supplements at different stages following differentiation protocols from previous reports^62^.

### Primary T cell cultures

After isolation, the CD4+ or CD8+ T Cells were cultured at a density of 0.5-1×10^6^/ml in complete XVivo15 medium (Lonza, 02-053Q) supplemented with 5% fetal bovine serum, 50 μM 2-mercaptoethanol (Sigma, M3148), 10 mM N-acetyl l-cysteine (Sigma, A7250), 1% penicillin/streptomycin (Life Technologies, 10378016). For CD4+ or CD8+ effector/memory differentiation, the medium was also supplemented with 10 ng/ml hIL-2 (R&D Systems, BT-002-050), 5 ng/ml of hIL-7 (R&D Systems, BT-007-AFL-025), 5 ng/ml hIL-15 (R&D Systems, BT-015-AFL-025)^63^. For CD4+ regulatory Th1 cell differentiation, the medium was supplemented with 10 ng/ml hIL-2, 5 ng/ml hIL-12 (R&D Systems, 10018-IL-010) and 1 ug/mL anti-IL4 (BioXcell, BE0240)^64^. T cells were activated with anti-CD3/CD28 Dynabeads at 1:1 ratio of the cells (ThermoFisher, 11131D) for 2 days in the complete XVivo15 medium with appropriate cytokines. After the activation, the Dynabeads were removed, the cells were edited by nucleofection and further cultured with the same complete medium with appropriate cytokines. Medium was changed every 2 days to keep the cell density below 1×10^6^/ml. The cells were cultured for 5 days (CD4+ and CD8+) or 7 days (Th1) before harvesting for STAG-seq.

### Cell lines

THP-1 and Jurkat cells were cultured in T75 flasks between 2-8x10^5^ cells/mL in RPMI1640 medium supplemented with 10% FBS (Sigma, F2442), 1% GlutaMAX (Life Technologies, 35050079), 1% penicillin/streptomycin (Life Technologies, 10378016).

### Flow-cytometry of HSPC-derived cells

At the indicated time points, cells were harvested by centrifugation at 400 x g for 5 minutes and washed twice with FACS buffer (PBS with 1% FBS and 2 mM EDTA). Cells were then stained with conjugated antibodies diluted in 100 uL FACS buffer for 30 minutes on ice while protected from light. After incubation, cells were washed twice and resuspended with the FACS buffer for flow cytometry analysis.

### In vitro transcription of base editor and prime editor mRNA

Base editor-encoded mRNA was transcribed in vitro with the protocol described previously ^66^. Briefly, the base editor cassettes (ORFs) were cloned into a plasmid containing an inactive T7 (dT7) promoter, 5′ untranslated region (UTR), Kozak sequence, and 3′ UTR (a gift from David Liu’s lab). All the components were PCR amplified with Q5® High-Fidelity 2X Master Mix (New England Biolabs, M0492L) using a forward primer that corrects T7 promoter sequence and a reverse primer that installed the poly(A) tail(119bp). The PCR products were gel purified and served as templates. mRNA was generated using the HiScribe T7 High-Yield RNA Kit (New England Biolabs, E2040S) according to the manufacturer’s instructions, with co-transcriptional capping by CleanCap AG (TriLink Biotechnologies) and full replacement of UTP with N1 Methylpseudouridine-50-triphosphate (TriLink Biotechnologies).

### Variant generation with guide RNA and Base editor mRNA electroporation

Cells were washed three times in DPBS with 0.1% BSA, and then resuspended in 10ul P3 Lonza buffer (Lonza, V4XP-3032). The editing master mix was prepared by combining 1.5ul base editor mRNA (2ug/ul) and 1.5uL of sgRNA (100 uM in IDTE pH 7.5, IDT), 10 ul P3 Lonza buffer. The cells were gently mixed with the master mix and transferred to the 20ul electroporation cuvette (Lonza, V4XP-3032). Immediately after electroporation, 120ul prewarmed appropriate medium was added to the cuvette. The cuvette was placed in a 37°C incubator for 20 min to allow the recovery of the cells.

Cells were then seeded in the appropriate complete medium.

### Gene knockout with RNP electroporation in T cells

T cells are prepared as above in 10ul P3 Lonza buffer. The editing master mix was prepared by combining 1.2uL of sgRNA (100 uM in IDTE pH 7.5, IDT) with 1ul S.p. HiFi Cas9 Nuclease (IDT, 1081061). The master mix was incubated at room temperature for at least 30 min, then diluted in 10 ul P3 Lonza buffer. The cells were then electroporated and recovered as above.

### Design of STAG-Seq Probes

The STAG-seq probes were designed as described previously^18^ To select the target sequences (52bp) of RNA, previously published guidelines were followed^18^.

### In house probe synthesis and preparation

For the 50K probe set, 5′ and 3′ probes were synthesized as separate oligonucleotide pools using Twist Bioscience’s oligo pool synthesis service. Oligo pools were diluted and pre-amplified by PCR, then transcribed the pool into RNA, followed by reverse transcription of the RNA back to single-strand probes.

### DNA amplicon panel design

The DNA panels were designed through Mission bio’s Tapestri Designer platform with GRCh38 genome. Each DNA panel contains the oligo pairs that generate 49 to 312 non-overlapping amplicons (175-275 bp) detecting the variant and control loci.

### STAG-seq Computational Pipeline

#### Quantification of RNA-transcript abundances

After demultiplexing the HyPR-Seq probe library, the raw .fastq files were processed into unique molecular identifier (UMI) count matrices using a modified HyPR-seq pipeline^18^, optimized for the STAG-seq read structure. The pipeline extracts the cell barcode from read 1 and the UMI + probe from read 2. The unique 3-tuples are then counted into an N_barcode x N_probe count matrix. In each cell, to compute RNA-transcript abundance for a specific gene, we sum the UMI counts for all probes targeting that gene. This produces a N_barcode x N_genes matrix where each entry represents the number of UMIs detected for a specific gene in a given cell.

#### Processing Genotype Information

After demultiplexing gDNA libraries, raw .fastq files were processed into .loom files using the Tapestri pipeline version 3 provided by MissionBio through their web portal. This pipeline requires both a reference genome and an amplicon panel file, which will be made available through the Gene Expression Omnibus (GEO) upon publication.

The Tapestri pipeline performs cell calling using a correlation UMAP algorithm to cluster valid and invalid cell barcodes. After cell calling, the pipeline genotypes individual cells with the Genome Analysis Toolkit (GATK).

#### Data Integration, Donor Separation, and Preprocessing

To integrate and process the data from HyPR-seq and Tapestri platforms, we employed custom Python scripts developed for STAG-Seq analysis. These scripts were built upon the Scanpy, AnnData, and h5pylibraries^68^.

#### Identifying Variants and Differential Expression Analysis

To classify genotyped variants within our dataset, cells exhibiting exactly one variant from our targeted set of variants were classified based on the genotype call provided by GATK, denoting them as either homozygous or heterozygous for that variant.

Additionally, cells were annotated as having a bystander mutation if another non-germline heterozygous or homozygous single nucleotide polymorphism (SNP) was detected within 10 base pairs of the targeted genomic edit.

